# Negative mood induction effects on problem solving task in women with eating disorders: a multi-method examination

**DOI:** 10.1101/467712

**Authors:** Melanie N. French, Kalina Eneva, Jean M. Arlt, Angelina Yiu, Eunice Y. Chen

**Author notes:** Corresponding author: Eunice Y. Chen, Ph.D., Temple Eating Disorders program, Temple University, 1701 North 13^th^ Street, Philadelphia, PA 19122. <u>Corresponding Author Contact information</u> **Email**.

## Abstract

Eating disorders (EDs) are a serious public health concern, affecting about 5.2% of American women. The effects of negative affect on problem-solving and its psychophysiological correlates are poorly understood in this population. This study examined respiratory sinus arrhythmia (RSA) and skin conductance responses of 102 women with EDs (Binge eating disorder [BED]: *n* = 57, Anorexia nervosa: *n* = 13, Bulimia nervosa [BN]: *n* = 32) and 24 healthy controls (HCs) at baseline, and then during: a negative mood induction task, an adapted Means Ends Problem-Solving (MEPS) task and recovery. The MEPS Task included four interpersonal scenarios: 1) binge-eating when sad, 2) job dissatisfaction, 3) feeling rejected by friends, and 4) jealousy in a relationship. ED groups reported more negative and less positive emotions than HCs. After a negative mood induction, women with BED provided significantly less effective problem-solving strategies compared to HCs and women with BN for the binge-eating MEPS scenario. Relative to baseline and the negative mood induction, all participants exhibited significantly higher skin conductance response and skin conductance levels throughout the MEPS scenarios and recovery. BED showed significantly lower RSA levels than individuals with BN and HCs throughout the protocol. The multimethod findings suggest individuals with BED are likely to solve problems by binge-eating when in a negative affective state.

## Introduction

Interpersonal problem-solving draws on both executive functioning abilities and emotion regulation, areas where eating disorder (ED) populations show particular weakness (Ivanova et al., 2015; Manasse et al., 2015). Although “top-down” emotion regulation strategies tend to be more effective than “bottom-up” strategies (McRae, Misra, Prasad, Pereira, & Gross, 2012), these pathways may compete during interpersonal problem-solving, and this competition may be stronger during periods of strong emotion. Respiratory sinus arrhythmia (RSA), a proximal measure of vagal tone captured through heart rate variability (HRV) over the respiration cycle (Beauchaine & Thayer, 2015; Berntson, Cacioppo, & Grossman, 2007), is a psychophysiological correlate of emotion regulation. In neutral states, the prefrontal cortex inhibits control of the amygdala and RSA increases (Peschel et al., 2016; Thayer, Ahs, Fredrikson, Sollers, & Wager, 2012); however, negative mood states place additional inhibitory challenges on this system, reducing RSA. Indeed, lower RSA has been observed across many clinical populations characterized by poor emotion regulation (Crowell, Beauchaine, & Linehan, 2009). After mood induction procedures, individuals with Anorexia-Nervosa (AN) and Binge Eating Disorder (BED; Friederich et al., 2006; Svaldi et al., 2016) showed reduced RSA levels compared to healthy participants. However, to date, there has not been an examination of a mood induction coupled with an interpersonal problem-solving task in ED populations.

The Means-Ends Problem Solving (MEPS) tests an established procedure that challenges individuals to generate solutions to realistic problem scenarios (Spivack, Platt, & Shure, 1976). Individuals with BED and subclinical disordered eating tend to generate less effective solutions on this task (Ridout, Matharu, Sanders, & Wallis, 2015; Svaldi, Caffier, & Tuschen-Caffier, 2010). Other studies using realistic problem-solving scenarios found that individuals with AN and bulimia nervosa (BN) performed worse on the Anorexia and Bulimia Problem Inventory, with particular weakness in weight and eating related scenarios (Espelage, Quittner, Sherman, & Thompson, 2000; Holt & Espelage, 2002). Though worsening problem-solving abilities following negative mood induction have been shown in other clinical samples, this has not been examined in an ED sample (Williams, Barnhofer, Crane, & Beck, 2005).

In our study, we used the MEPS task and psychophysiological measures to examine problem-solving after a mood induction procedure in women with AN, BN, BED, and healthy control (HC) women. We elicited individuals’ autobiographical memories for the mood induction to elicit “top-down” emotion regulation strategies (McRae et al., 2012) and to facilitate pathway competition during the problem-solving task. This technique has successfully induced negative mood in other studies using ED samples (Hilbert, Vogele, Tuschen-Caffier, & Hartmann, 2011; Telch & Agras, 1996). We measured skin conduction level as a manipulation check for physiological arousal (Boucsein, 1992), during the mood induction and the MEPS procedure. In addition, because the MEPS does not include disorder-specific scenarios, and previous research has shown that individuals with EDs have poorer performance in these domains (Espelage et al., 2000; Holt & Espelage, 2002), we included a binge-eating scenario during the task. A binge-eating scenario was chosen above a restrictive eating behavior or compensatory behavior because urges to binge eat occur across all eating disorder subtype groups (Hawkins & Clement, 1984).

We hypothesized that women diagnosed with AN, BN, and BED, compared to HC women, (1) would have greater self-reported negative affect and less self-reported positive affect, and higher urges to binge-eat in response to a negative mood induction and throughout the protocol. We also hypothesized that (2) women with EDs would generate fewer and less effective solutions compared to HCs, especially on the binge-eating scenario, and that (3) during the negative mood induction and throughout the MEPS task, individuals with EDs would show lower RSA compared to HCs. Finally, as a manipulation check, we predict that (4) higher SCRs and tonic SCL values during the mood induction and MEPS for all participants.

## Methods

### Participants

The sample consisted of 102 women who sought treatment at an adult Eating and Weight Disorders Clinic and 24 participants without a history of EDs or other Axis I or II disorders per the Diagnostic Statistical Manual-IV-Text Revision (Association, 2000). Our study was carried out in accordance with The Code of Ethics of the Declaration of Helsinki (Rits, 1964). All participants gave written consent to participate in this study. Of the participants who met DSM-IV-TR criteria for an ED, 57 met criteria for BED, 13 for either AN or sub-clinical AN, and 32 for BN (Association, 2000). Only women were recruited due to indicated differences in emotional reactions (Fischer, 2000) as well as differences in cardiovascular control and vagal activity between genders (Ryan, Goldberger, Pincus, Mietus, & Lipsitz, 1994). Participants were also excluded if they were taking tranquilizers, antihistamines, scopolamine, or beta blockers due to their enhancement of parasympathetic nervous system activity (Elenkov, Wilder, Chrousos, & Vizi, 2000). Due to the wide use of antidepressants in the treatment of EDs, we did not exclude individuals with EDs who were taking antidepressants. However, individuals with EDs who were on selective serotonin reuptake inhibitors were excluded if they had not been stable on the same antidepressant and dosage for the last 3 months.

### Measures

#### Eating disorder symptoms

The Eating Disorders Examination (EDE) Version 16.0 (Fairburn, Cooper, & O’Conner, 2008) is a standardized semi-structured interview, measuring the frequency and severity of ED psychopathology and key behaviors. The EDE has good internal consistency, high test-retest reliability, and inter-rater reliability (Rizvi, Peterson, Crow, & Agras, 2000). The EDE yields subscales of restraint, eating concerns, shape concerns, and weight concerns.

#### Psychiatric conditions

The Structured Clinical Interview for DSM–IV–TR (SCID-(First, Spitzer, Gibbon, & Williams, 2002) is a semi-structured clinical interview used to assess for Axis I disorders per the DSM-IV-TR (Association, 2000). The SCID-I has adequate interrater reliability (First et al., 2002).

#### State measures

Participants reported their emotional states using the Visual Analogue Scale (VAS) (Haines, Williams, Brain, & Wilson, 1995) three times: (1) before, (2) after negative mood induction, and (3) after the MEPS. See Figure 1. The VAS is a state measure of emotions on a continuous scale ranging from 0 to 100 on the following emotions: frustration, anxiety, happiness, tension, fear, sadness, and “urge to binge”. Higher scores indicate a greater experience of the emotion.

**Figure 1:**
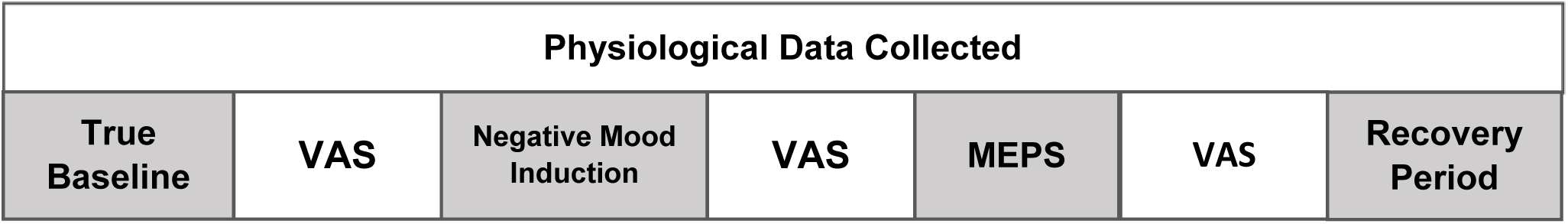
Timeline of Procedure starting from baseline to the Means-End Problem Solving (MEPS) to the recovery period with visual analogue scales (VAS) of emotion taken in between time points.

#### Sad memories questionnaire

As per the procedure described by Hernandez, Vander Wal, and Spring (2003), participants were asked to generate one-sentence descriptions of ten sad autobiographical memories that were rated on 10-point Likert scales to reflect (a) how sad the memory made them feel, and (b) how vividly they recalled the memory. The three memories with the highest averaged vividness and intensity ratings were chosen for the negative mood induction.

### Procedures

#### Negative mood induction

Please see Figure 1. Negative mood was induced through autobiographical memories and sad music, as per the protocol of Hernandez, Vander Wal, and Spring (2003). During an 8 minute negative mood induction, each individual listened on earphones to a compilation of classical music that has been shown to induce sad moods (i.e., unpleasant affect and low arousal while being instructed to recall their preselected autobiographical memories (Eich, Macaulay, & Ryan, 1994).

#### Behavioral measures of problem-solving

The Means-Ends Problem-Solving Task (MEPS; Spivack et al., 1976) was used to assess problem-solving, where the participants were provided with a scenario that describes a problem and a stated outcome. The participant was instructed to generate the steps, or means, that could be taken to achieve the stated outcome. In the current study, four sets of MEPS items were adapted from the original task (Spivack et al., 1976).The scenarios consisted of problems surrounding binge-eating, job performance, and two interpersonal scenarios (one between friends and one with a significant other). The ‘Bingeeating’ scenario was a new scenario developed for the ED population, which presented the story of an individual who considered binge-eating when feeling sad, and it ended with her not engaging in a binge. The ‘Performance’ scenario introduced a situation where an individual felt dissatisfied with her job responsibilities and was considering a new career, and it finished with her satisfied in her career. The ‘Friends’ scenario described an individual who felt as if her friends have been avoiding her, and it resolved with them all getting along. The ‘Significant Other’ scenario presented an individual whose significant other was flirting with another person and concludes with the two reconciling.

The adapted MEPS task (Williams et al., 2005) gave participants 60 seconds to describe the most effective strategy for solving the problem and additional 60 seconds to describe alternative strategies. For each of the four MEPS scenarios, two dependent variables were derived: the overall effectiveness of the participant’s response, which was blindly rated on a 7-point scale (1 = not at all effective to 7 = extremely effective), and the number of relevant means (active problem-solving steps) the participant produced. The MEPS task was recorded, transcribed, and scored by an independent rater who was blind to group status.

### Physiological measures

#### Respiratory sinus arrhythmia

RSA was assessed during 4 different phases: (1) 5-minute baseline, (2) 8-minute negative mood induction, (3) 20-minute MEPS task, and then (4) during a 5-minute recovery period. We collected electrocardiogram (ECG) data utilizing a modified Lead II configuration (Mindware Technology LTD; Gahanna, Ohio US, 43230) and respiration response using a chest respiration belt (AMBU Sleepmate; Ballerup Denmark). At baseline, the participant was instructed to ‘relax without focusing on anything in particular’. See Figure 1.

RSA was used as the index of vagal tone and was measured by assessing the high frequency band of spectral analysis (Berntson, Cacioppo, & Fieldstone, 1996), which decomposes electrocardiogram (ECG) R-wave time series into three HRV frequency ranges through Fast Fourier transformations. Spectral analyses were conducted using Mindware Technologies HRV 2.33 software (Westerville, OH), which detects questionable RR intervals on the basis of the overall RR distribution using a validated algorithm to aid artifact detection and editing (Berntson, Quigley, Jang, & Boysen, 1990). The low frequency range is less than 0.04 Hz, the mid-frequency ranges from 0.04 Hz to 0.15 Hz, and the high frequency range is greater than 0.15 Hz. Research on heart rate and HRV suggests that parasympathetic/vagal activity influences all frequencies of <0.5, whereas sympathetic activity affects frequencies of <0.15 (Berntson et al., 1996). High frequency spectral densities were calculated across 60-second intervals.

#### Skin conductance responses

Like RSA, SCRs were assessed during the same 4 different phases (Figure 1). Skin conductance responses (SCRs) and skin conductance level (SCL) were used as measures of sympathetic activity (Boucsein, 2012a). Skin conductance was recorded from two electrodes attached to the palm of the non-dominant hand using a galvanic skin conductance device made by Mindware Technology LTD; Gahanna, Ohio USA 43230. Increases in skin conductance have been associated with negative affect following exposure to emotional stimuli, even when cardiac measures did not change significantly (Salters-Pedneault, Gentes, & Roemer, 2007). SCRs were used, as they are a preferred electrodermal measure for tasks eliciting emotional activation (Boucsein, 2012b). Tonic SCL referred to the level of skin conductance per data collection period.

### Statistical treatment

Preliminary analyses included an assessment of group differences in demographic variables, medical, and psychiatric co-morbidities using chi-square tests and ANOVAs.

Three repeated measures using Time (baseline, during the mood induction, and during the MEPS) x Group (BN, BED, AN, and HC) ANOVAs were conducted to assess self-reported negative emotions, happiness, and urges to binge-eat.

For each of the four MEPS scenarios, a univariate ANOVA was used to examine group differences in the number of relevant means and effectiveness of solutions for each of the four MEPS scenarios.

Three separate repeated measures using Time (baseline, mood induction, each MEPS scenario, recovery) x Group (BN, BED, AN, HC) ANOVAs were conducted for RSA, SCR, and tonic SCL. Partial η2 was used to report effect sizes for repeated measures ANOVAs and ANOVAs, with the following cut-off conventions: small (.01), medium (.06) and large (.14).

## Results

### Sample description

Please see Table 1 for a description of the means, standard deviations and group sociodemographic differences of the sample. There were no significant group differences on race, marital status, or full-time employment/student status. There were significant groups differences on age and annual income below $25,000 (i.e. low-income status), such that individuals with BED were older relative to individuals with BN, AN, and HCs, and were more likely to earn an annual income above $25,000. However, older age and higher income were correlated in the sample (r = .349; p<.001).

**Table 1:**
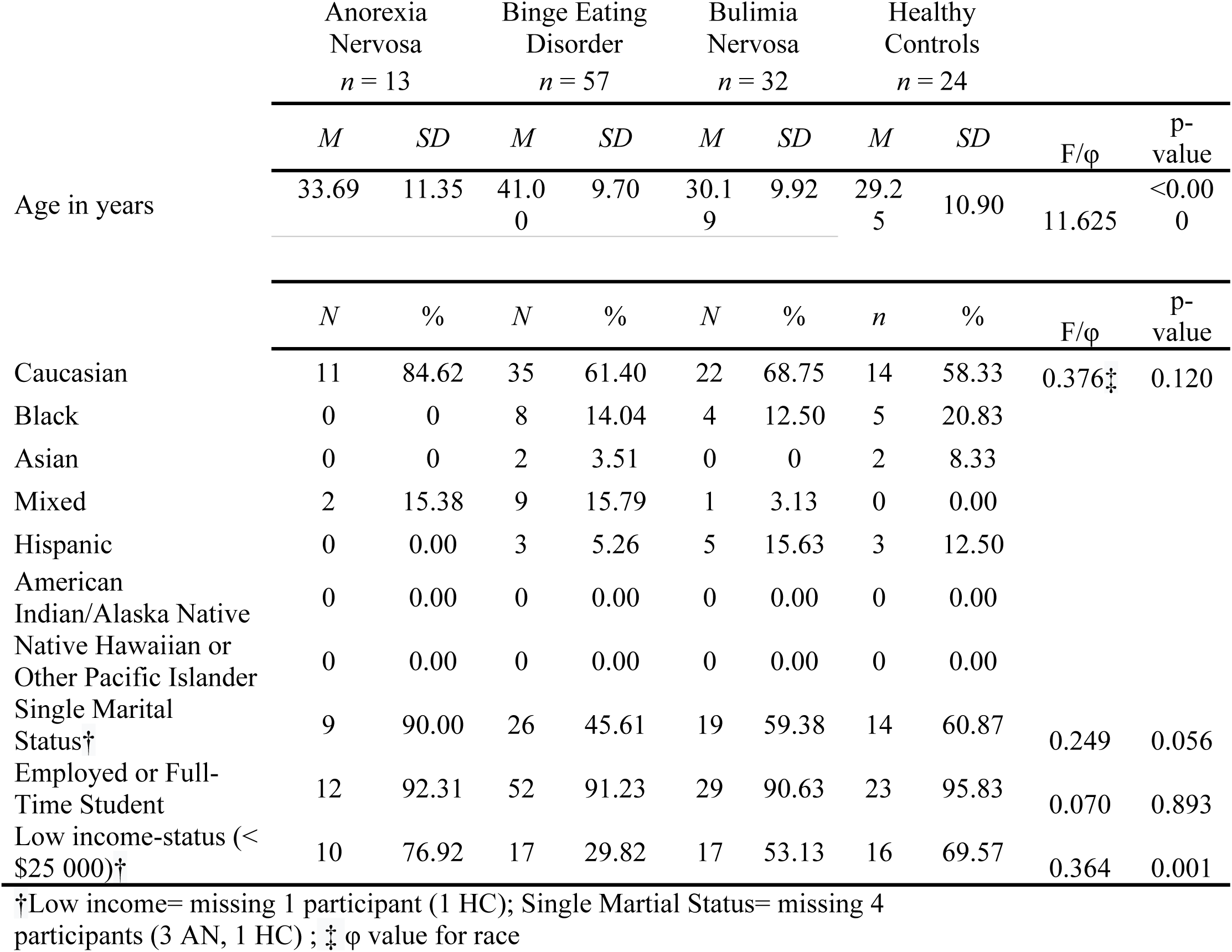
Demographic information of sample (means and standard deviations).

Table 2 describes the clinical characteristics of the sample. Overall, group differences in clinical characteristics were aligned with diagnosis status. As expected, the BED group had the highest BMI, followed by HCs and the BN group, and, lastly, the AN group had the lowest BMI (all group differences were significant except the BN vs HCs; *p*’s<.001). ED groups had greater eating disorder psychopathology (as measured by the EDE subscales) and higher lifetime prevalence mood and anxiety disorders compared to HCs. Additionally, the AN group had more anxiety disorders than the BED group, and the BN and AN groups scored higher on the EDE dietary restraint scale than the BED groups. Furthermore, the BN and BED groups had a higher rate of medication use and medical problems (such as high blood pressure, asthma, high cholesterol, etc.) compared to HCs, and the BN group also had more medical problems than the BED group. Consistent with clinical presentation, and the BED and BN groups engaged in more bingeing than the AN and HC groups. Finally, those with BN had the highest frequency of vomiting episodes than all other groups.

**Table 2:** Clinical characteristics of the sample (means and standard deviations).

|  | Anorexia Nervosa<br>*n*=13 |  | Binge Eating Disorder<br>*n* = 57 |  | Bulimia Nervosa<br>*n* = 32 |  | Healthy Controls<br>*n* = 24 |  |  |  |
| --- | --- | --- | --- | --- | --- | --- | --- | --- | --- | --- |
|  | *M* | *SD* | *M* | *SD* | *M* | *SD* | *M* | *SD* | F/φ | p-value |
| BMI ( kg/m2) | 18.83 | 3.82 | 36.55 | 10.21 | 26.13 | 6.03 | 26.92 | 4.63 | 25.42 | <0.000 |
| EDE-EC | 3.52 | 1.76 | 2.82 | 1.40 | 2.74 | 1.20 | 0.04 | 0.08 | 34.68 | <0.000 |
| EDE-DR | 3.65 | 1.74 | 2.13 | 1.27 | 3.14 | 1.32 | 0.52 | 0.92 | 25.12 | <0.000 |
| EDE-WC | 3.12 | 1.70 | 3.57 | 1.08 | 3.45 | 1.17 | 0.52 | 0.69 | 45.68 | <0.000 |
| EDE-SC | 3.75 | 1.45 | 3.82 | 1.28 | 3.72 | 1.09 | 0.60 | 0.81 | 47.36 | <0.000 |
| Objective Binge Episodes | 1.31 | 2.59 | 4.49 | 4.09 | 6.55 | 6.43 | 0.00 | 0.00 | 12.30 | <0.000 |
| Vomit Episodes | 2.09 | 3.93 | 0.13 | 0.93 | 7.34 | 8.75 | 0.00 | 0.00 | 18.81 | <0.000 |
|  | *N* | % | *N* | % | *N* | % | *N* | % | F/φ | p-value |
| Mood Disorders | 7 | 53.85 | 37 | 64.91 | 24 | 75.00 | 0 | 0.00 | 0.54 | <0.000 |
| Anxiety Disorders | 9 | 69.23 | 21 | 36.84 | 12 | 37.50 | 0 | 0.00 | 0.40 | <0.000 |
| Medical Comorbidities† | 7 | 63.64 | 27 | 50.94 | 21 | 75.00 | 4 | 21.05 | 0.35 | 0.003 |
| Medication Use | 6 | 46.15 | 29 | 50.88 | 15 | 46.88 | 2 | 8.33 | 0.33 | 0.004 |

ABBREVIATIONS: Body Mass Index (BMI); Dietary Restraint (DR); Eating Disorder Examination (EDE); Eating Concerns (EC); Shape Concerns (SC); Weight Concerns (WC)
†Medical Comorbidity: missing 15 participants (2 AN, 4 BED, 4 BN, & 5 HC);
Note: All eating behavior is within the previous three months' time frame

### Main analyses

#### Hypothesis 1: Self-reported emotions

Overall, there was a main effect of Time on emotions, such that all participants reported more negative emotions, *F*_(2, 107)_ = 28.98, *p* < .001, η^2^ =.21, and fewer positive emotions, *F*_(2,107)_ = 56.88, *p* < .001, η^2^ =.35, after completion of the negative mood induction, in comparison to baseline and after the MEPS task. There was a main effect of Group on negative emotions (*F*_(3,107)_ = 4.31, *p* = .007, η^2^ =.11) and positive emotions (*F*_(3, 107)_ = 5.49, *p* = .002, η^2^ =.13). Individuals with EDs reported less positive emotions than HC participants. Participants with BN and BED reported more negative emotions than HCs, and there was a trend for participants with AN to report more negative emotions than HCs (*p* =.097). There was no Time × Group interaction for negative emotions or positive emotions. For urges to engage in binge-eating, there was a main effect of Group, *F*_(3, 107)_ = 10.12, *p* < .001, η^2^ =.22, such that individuals with BED and BN demonstrated significantly greater urges to binge-eat compared to the HC subgroup (*p*’s < .001), although there was no effect of Time (*p* = .926) or Time × Group interaction *(p* = .426). See Table 3 for group means and standard deviations.

**Table 3:** Mean and standard deviations of self-reported negative emotions, positive emotions and urges to binge-eat for groups prior to the Negative Mood Induction, after the Negative Mood Induction, and after the Means-End Problem Solving (MEPS) task.

|  | Negative Emotions Prior<br>to Mood Induction | Negative Emotions<br>After Mood Induction | Negative Emotions<br>After MEPS |
| --- | --- | --- | --- |
|  | *M (SD)* | *M (SD)* | *M (SD)* |
| Anorexia Nervosa (*n* = 9) | 32.49 (26.67) | 42.76 (23.27) | 44.49 (23.94) |
| Binge Eating Disorder (*n* = 55) | 26.73 (20.06) | 47.99 (27.366) | 33.17 (23.22) |
| Bulimia Nervosa (*n* = 28) | 26.73 (21.69) | 62.20 (18.90) | 35.38 (22.56) |
| Healthy Control (*n* = 19) | 11.87 (16.43) | 38.44 (30.62) | 20.31 (21.85) |
|  | Positive Emotions<br>Prior to Mood Induction | Positive Emotions<br>After Mood Induction | Positive Emotions<br>After MEPS |
|  | *M (SD)* | *M (SD)* | *M (SD)* |
| Anorexia Nervosa (*n* = 9) | 31.02 (22.05) | 3.44 (4.35) | 20.94 (22.38) |
| Binge Eating Disorder (*n* = 55) | 36.27 (23.28) | 10.41 (18.34) | 28.07 (19.85) |
| Bulimia Nervosa (*n* = 28) | 31.60 (24.65) | 7.12 (14.69) | 23.89 (23.31) |
| Healthy Control (*n* = 19) | 53.72 (21.58) | 20.65 (22.92) | 43.52 (28.78) |
|  | Urge to Binge<br>Prior to Mood Induction | Urge to Binge<br>After Mood Induction | Urge to Binge<br>After MEPS |
|  | *M (SD)* | *M (SD)* | *M (SD)* |
| Anorexia Nervosa (*n* = 9) | 29.78 (36.68) | 16.65 (33.35) | 23.78 (36.21) |
| Binge Eating Disorder (*n* = 55) | 33.70 (30.87) | 40.98 (35.30) | 36.08 (31.06) |
| Bulimia Nervosa (*n* = 28) | 26.98 (27.04) | 31.88 (35.34) | 34.70 (36.46) |
| Healthy Control (*n* = 19) | 0.65 (1.81) | 0.51 (1.05) | 0.43 (0.95) |

*Note:* Self-reported ratings were made on a 0-100 Visual Analog Scale; All participants missing visual analog data points were excluded from the analysis.

#### Hypothesis 2: MEPS generated solutions and effectiveness

A univariate ANOVA suggests that there were significant group differences between effectiveness of the solutions generated for the ‘Binge-Eating’ scenario, *F*_(3,93)_ = 5.960, *p* = .001. Women with BED generated significantly less effective solutions than HCs and women with BN (*p* <.001 and *p* =.002 respectively), although there were no differences in the number of relevant solutions generated. There were no differences between diagnostic groups on the remaining three MEPS scenarios (*p*s > .05). See Table 4 for group means and standard deviations.

**Table 4:** Mean and standard deviations of relevant and effective solutions generated in the Means-End Problem Solving (MEPS) for Bulimia Nervosa, Binge Eating Disorder, Anorexia Nervosa, and Healthy Controls.

| **Binge Eating Scenario** | Relevant Solutions<br>*M (SD)* | Effectiveness<br>*M (SD)* |
| --- | --- | --- |
| Anorexia Nervosa | 7.75 (3.37) | 3.75 (1.91) |
| Binge Eating Disorder | 5.79 (2.44) | 3.44 (1.48) |
| Bulimia Nervosa | 6.40 (2.75) | 4.57 (1.48) |
| Healthy Control | 6.80 (2.44) | 4.95 (1.28) |
| **Job Performance Scenario** | Relevant Solutions<br>*M (SD)* | Effectiveness<br>*M (SD)* |
| Anorexia Nervosa | 5.25 (2.55) | 4.50(1.69) |
| Binge Eating Disorder | 5.26 (2.39) | 3.56 (1.52) |
| Bulimia Nervosa | 4.60 (1.67) | 4.03 (1.35) |
| Healthy Control | 4.95 (1.85) | 4.15 (0.93) |
| **Friends Scenario** | Relevant Solutions<br>*M (SD)* | Effectiveness<br>*M (SD)* |
| Anorexia Nervosa | 3.88 (1.25) | 3.38 (1.77) |
| Binge Eating Disorder | 4.00 (1.41) | 3.21 (1.03) |
| Bulimia Nervosa | 4.37 (2.22) | 3.47 (1.55) |
| Healthy Control | 4.75 (1.59) | 4.00 (1.12) |
| **Significant Other Scenario** | Relevant Solutions<br>*M (SD)* | Effectiveness<br>*M (SD)* |
| Anorexia Nervosa | 3.88 (1.13) | 2.88 (1.13) |
| Binge Eating Disorder | 4.69 (1.98) | 3.62 (1.55) |
| Bulimia Nervosa | 4.07 (2.07) | 3.27 (1.55) |
| Binge Eating Disorder | 4.69 (1.98) | 3.62 (1.55) |

Notes: Number of relevant means had no upper end; Effectiveness was scored on a scale of 1-7; All participants missing visual analog data points were excluded from the analysis.

#### Hypothesis 3: Respiratory sinus arrhythmia

There was a significant effect of Group for RSA values (*F*_(3, 122)_ = 5.60, *p* = .001, η^2^ = .12), such that individuals with BED exhibited significantly lower RSA levels than individuals with BN (*p* =.001) and HCs (*p* = .002) throughout the protocol. There was not a significant effect for Time (*p* =.312) or for the Time X Group interaction (*p* =.211). See Figure 2 and Supplemental Table 5 for RSA findings.

**Figure 2:**
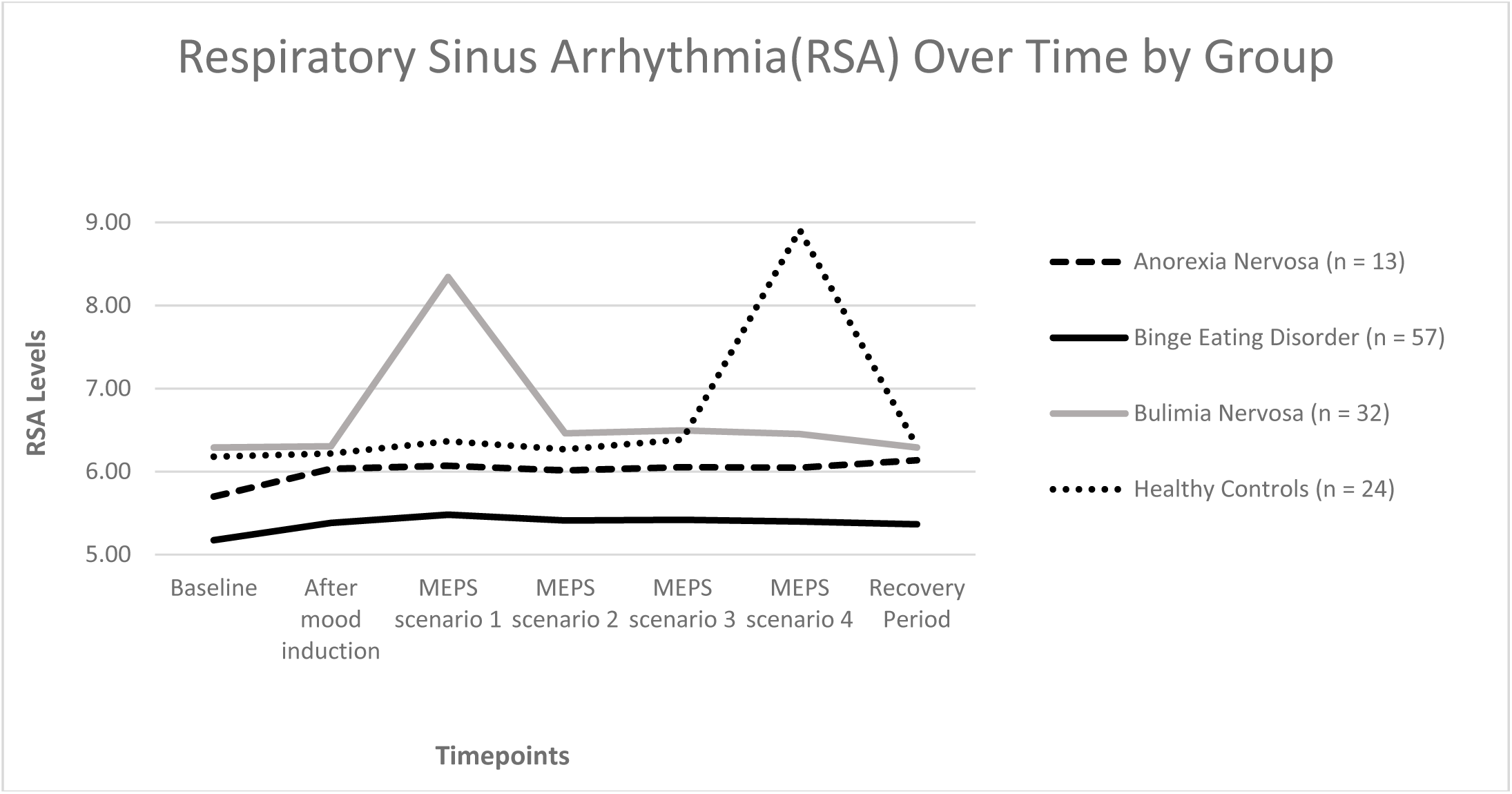
Means of Respiratory Sinus Arrhythmia Values at: 1) Baseline, 2) Negative Mood Induction, 3) Means-End Problem Solving (MEPS) Scenario 1: Binge eating, 4) MEPS Scenario 2: Job Performance, 5) MEPS Scenario 3: Friends, 6) MEPS Scenario 4: Significant other, and 7) Recovery

#### 3.2.4 Hypothesis 4: Skin conductance

There was a significant effect of Time for SCRs (*F*_(6,89)_ = 40.86, *p* < .001, η^2^ = .32) and tonic SCL values (*F*_(6, 92)_ = 46.83, *p* < .001, η^2^ = .34). Relative to baseline and the negative mood induction, all participants exhibited significantly higher SCR and SCL levels throughout the MEPS scenarios and during the recovery period (*p*’s <.001). For SCR and tonic SCL values, there were no significant effects of Group (*p* =.352, *p* = .943, respectively) or Time X Group interaction (*p* =.462, *p* = .441). SCR and SCL are depicted in Figure 3 and 4 respectively (see also Supplemental Tables 6 and 7 respectively).

**Figure 3:**
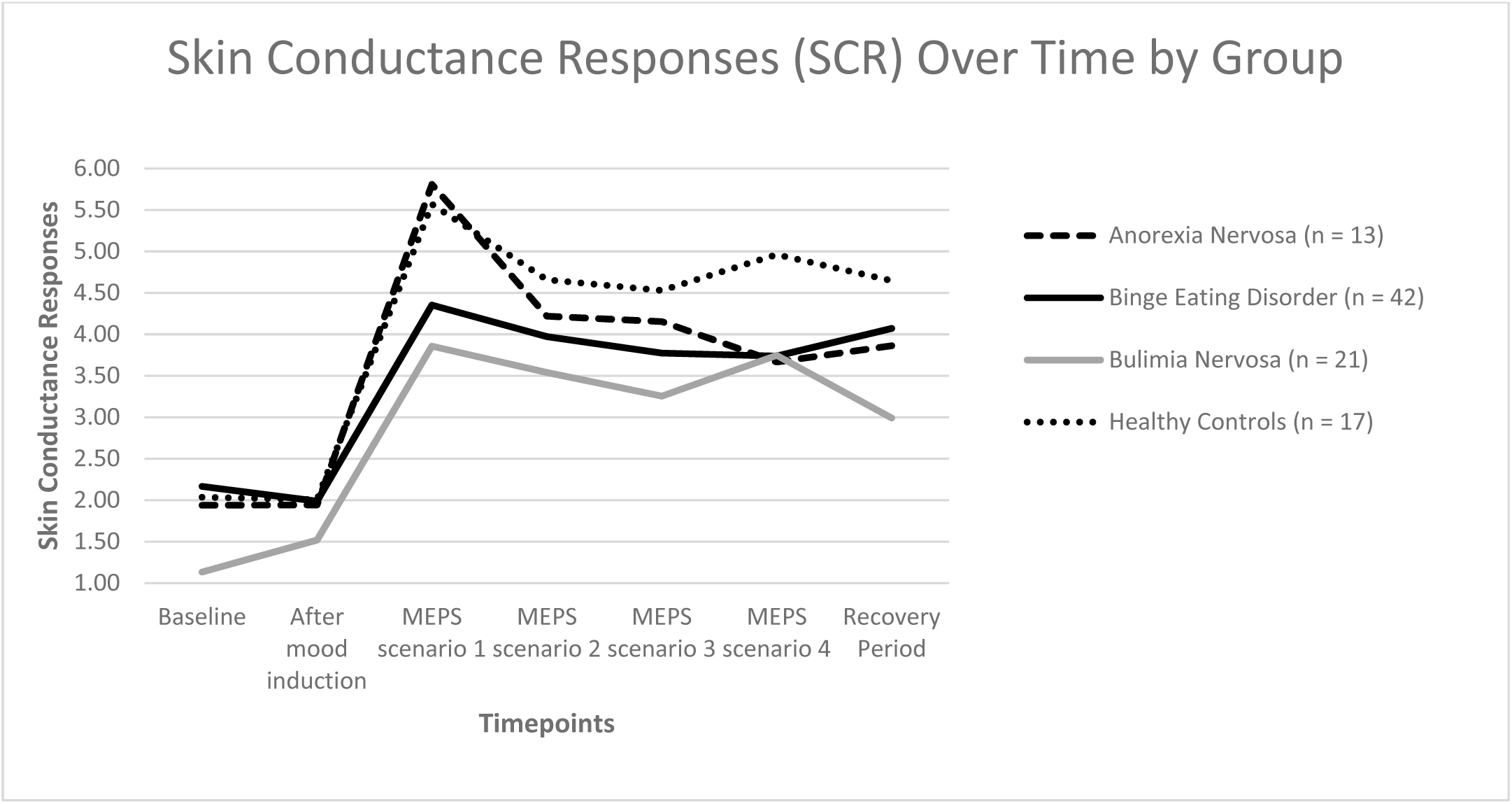
Means of Skin Conductance Responses at: 1) Baseline, 2) Negative Mood Induction, 3) Means-End Problem Solving (MEPS) Scenario 1: Binge eating, 4) MEPS Scenario 2: Job Performance, 5) MEPS Scenario 3: Friends, 6) MEPS Scenario 4: Significant other, and 7) Recovery. Participants were excluded from the analysis with missing skin conductance responses values.

**Figure 4:**
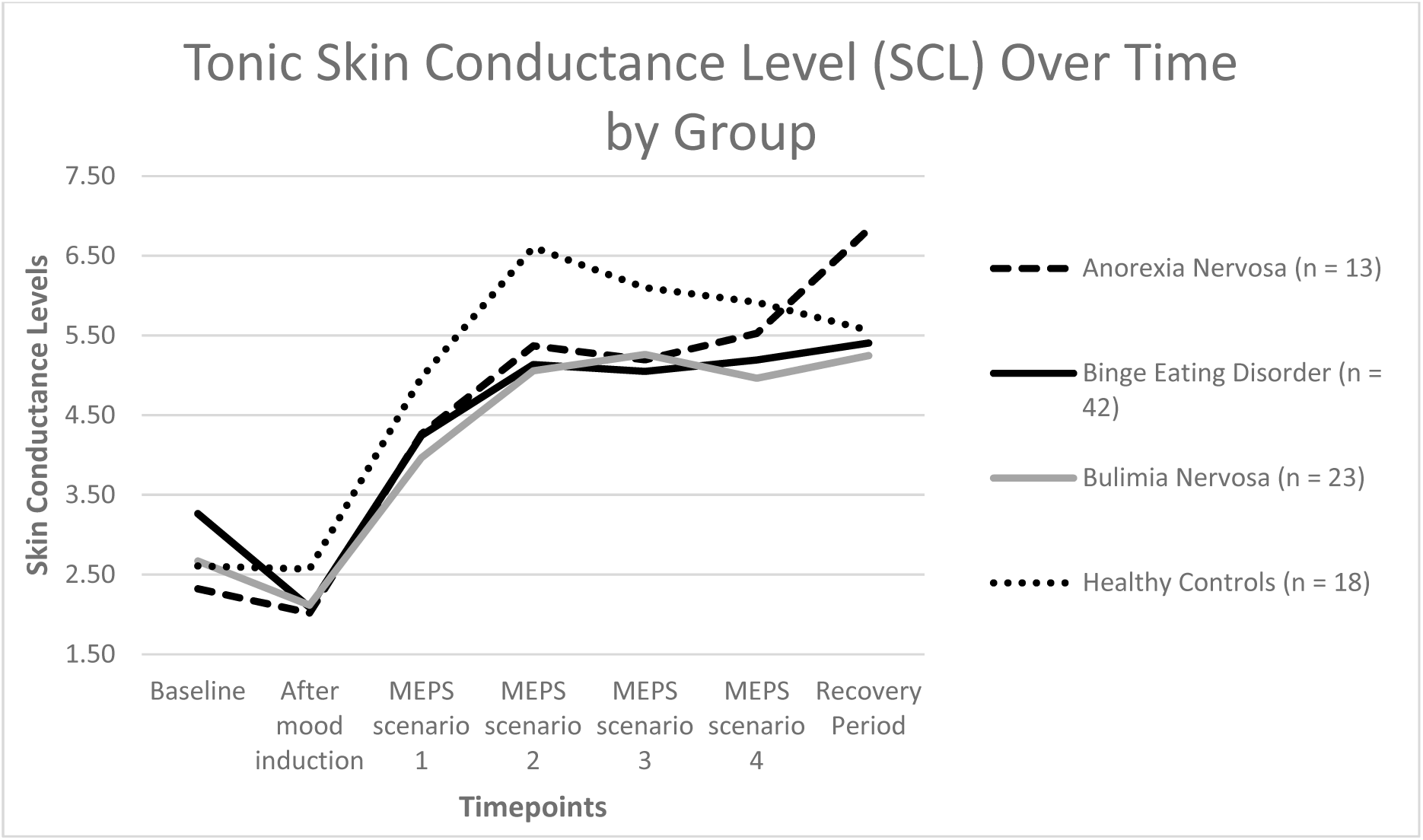
Means of Skin Conductance Levels at: 1) Baseline, 2) Negative Mood Induction, 3) Means-End Problem Solving (MEPS) Scenario 1: Binge eating, 4) MEPS Scenario 2: Job Performance, 5) MEPS Scenario 3: Friends, 6) MEPS Scenario 4: Significant other, and 7) Recovery. Participants were excluded from the analysis with missing skin conductance responses values.

## Discussion

This study used a multi-method design to test the effects of a negative mood induction on interpersonal problem-solving, self-reported emotions, and psychophysiological measures. The mood induction component of the study was effective, which conforms with previous studies using clinical and subclinical ED samples (Cardi, Leppanen, & Treasure, 2015; Chua, Touyz, & Hill, 2004; Rommel et al., 2015; Sheppard-Sawyer, McNally, & Fischer, 2000; Svaldi et al., 2010), including those using autobiographical techniques (Svaldi et al., 2016; Telch & Agras, 1996). Self-reported negative emotions increased following a negative mood induction across a sample of women with EDs and HCs. Concordantly, a psychophysiological manipulation check revealed, after the mood induction, SCL and SCL increased across all groups. Additionally, on the binge-eating scenario from the MEPS, the BED sample generated significantly fewer effective solutions than HCs and the BN group. Finally, individuals with BED group significantly lower RSA than individuals with BN and HCs throughout the protocol.

### Problem-solving challenges after a “top-down” mood induction

Our study examines “top-down” problem-solving after a mood induction procedure that were used to facilitate pathway competition. We found that during a negative mood induction, while completing an interpersonal problem-solving task, women with BED generated less effective solutions with a disorder-specific problem-solving scenario relative to HCs. A systematic review by Kittel, Brauhardt, and Hilbert (2015) revealed that women with BED perform slightly worse in certain domains of cognitive functioning compared to normal weight and overweight HCs, and this performance worsens with disorder-related stimuli (e.g. food) Additionally, Danner et al. (2013) found that negative affect hinders decision-making in individuals with BED relative to HCs. Extending previous findings, our study suggests that deficits in cognitive functioning after a negative affect induction are particularly pronounced for disorder-specific stimuli in BED.

We did not observe poorer performance in the other MEPs interpersonal problem-solving scenarios in individuals with BED. Previous studies suggests that individuals with BED often have problems in social and interpersonal domains (Aloi, Rania, Caroleo, De Fazio, & Segura-García, 2017; Blomquist, Ansell, White, Masheb, & Grilo, 2012; Tasca, Balfour, Presniak, & Bissada, 2012), especially during negative affect (Ivanova et al., 2015), and one previous study using a shortened MEPS task reported individuals with BED produced less effective solutions on interpersonal scenarios compared to overweight HCs (Svaldi, Dorn, & Trentowska, 2011). More research is needed to confirm if challenges in problem solving are disorder-specific or more general across interpersonal situations and whether these difficulties are particularly enhanced with a negative mood induction.

Additionally, we found that the BED group generated less effective solutions than the BN group in the binge eating scenario as well. As the BED group had a significantly higher BMI than the BN group, it may be that obesity compounds the difficulties in problem-solving under stress. Obesity in and of itself has shown to negatively influence certain domains of executive functioning (Yang, Shields, Guo, & Liu, 2018). This interaction between BED-related pathologies (i.e. reduced cognitive resources when under stress) and obesity may increase challenges in problem-solving tasks.

### Mood induction and psychophysiological results

The BED group exhibited lower RSA compared to the BN and HC groups. Lower RSA is associated with a wide range of psychological problems, including emotional rigidity and poor social functioning (Carney et al., 2001). Research suggests that individuals with BED have poorer emotion regulation skills than HC groups (Gianini, White, & Masheb, 2013; Kenardy, Arnow, & Agras, 1996). Past psychophysiological experiments using individuals with BED also reported a decreased parasympathetic response to stress (Friederich et al., 2006; Svaldi et al., 2010), although one other study did not show consistent psychophysiological differences (Hilbert et al., 2011), suggesting more studies need to be done to verify our findings. However, our findings fit with a model of reduced RSA as being associated with negative mood and stress on the parasympathetic system (Thayer et al., 2012).

As expected SCR and SCL increased for all groups after the mood induction, during the MEPs scenarios and during recovery. In the literature, there have been inconsistent effects of negative mood induction and/or cognitive on skin conductance levels. Some studies using ED samples showed that SCL increased after a negative mood induction or a cognitive task (Tuschen-Caffier & Vogele, 1999). After a negative mood induction, another study found no change in SCL (Svaldi et al., 2010), and another study found that SCL decreased (Hilbert et al., 2011). The mix of findings suggest the need for further studies examining the effect of negative mood induction and binge eating scenarios on skin conductance measures.

### Strengths and limitations

Our study did not include separate control groups of overweight and normal-weight individuals without EDs in order to dissociate the effects of weight. As weight can influence RSA, this may account for the few psychophysiological differences observed between the ED and HC groups. Additionally, high BMI or other coexisting-medical conditions (such as diabetes and hypertension) may cause blunted cardiovascular reactivity (Carroll, Phillips, Der, Hunt, & Benzeval, 2011; Masi, Hawkley, Rickett, & Cacioppo, 2007), which may have lowered RSA levels in the BED group. However, other studies controlling for weight have shown that women with EDs have challenges in emotion regulation (Kittel et al., 2015; Svaldi et al., 2010; Telch & Agras, 1996) and problem-solving (Eneva, Arlt, Yiu, Murray, & Chen, 2017; Kittel et al., 2015; Manasse et al., 2015) domains. In addition, we were also underpowered to detect significant effects for individuals with AN, and would have benefited from additional participants in this group. Our study did not compare MEPs performance before and after a negative mood induction. The order of MEPS scenarios between the binge-eating and general scenarios was also not counterbalanced.

Study strengths included the use of a disorder-specific problem-solving scenario in a heterogeneous, clinical ED sample after a mood induction procedure and used a multi-modal data collection approach.

### Conclusion & future directions

Our results show significant differences in “top-down” emotional regulation during interpersonal problem-solving in individuals with BED using self-report, behavioral, and psychophysiological data. Individuals with BED may have a cognitive vulnerability to negative affect that may lead to poorer interpersonal problem-solving related to disorder-specific scenarios (i.e. binge-eating). Use of novel approaches to assess ambulatory responses to negative affect induction and disorder-specific problems to test a ‘top-down’ mood induction problem-solving model in real-life are needed in the future.

## Acknowledgements

This work was supported by the National Institute of Mental Health [grant numbers R21MH093932-01A1;K23MH081030-01A1].

